# Neural divergence and hybrid disruption between ecologically isolated *Heliconius* butterflies

**DOI:** 10.1101/2020.07.01.182337

**Authors:** Stephen H. Montgomery, Matteo Rossi, W. Owen McMillan, Richard M. Merrill

## Abstract

The importance of behavioural evolution during speciation is well established, but we know little about how this is manifest in sensory and neural systems. Although a handful of studies have linked specific neural changes to divergence in host or mate preferences associated with speciation, how brains respond to broad environmental transitions, and whether this contributes to reproductive isolation, remains unknown. Here, we examine divergence in brain morphology and neural gene expression between closely related, but ecologically distinct, *Heliconius* butterflies. Despite on-going gene flow, sympatric species pairs within the *melpomene-cydno* complex are consistently separated across a gradient of open to closed forest and decreasing light intensity. By generating quantitative neuroanatomical data for 107 butterflies, we show that *H. melpomene* and *H. cydno* have substantial shifts in brain morphology across their geographic range, with divergent structures clustered in the visual system. These neuroanatomical differences are mirrored by extensive divergence in neural gene expression. Differences in both morphology and gene expression are heritable, exceed expected rates of neutral divergence, and result in intermediate traits in first generation hybrid offspring. This likely disrupts neural system function, leading to a mismatch between the environment and the behavioral response of hybrids. Our results suggest that disruptive selection on both neural function and external morphology result in coincident barriers to gene flow, thereby facilitating speciation.

## Introduction

Ecological adaptation is a major force driving the evolution of new species [1,2]. Although it is well established that divergent selection can influence behavioural traits and promote speciation [3], there are few empirical examples of how divergent selection acts on the underlying sensory and neural systems. For example, existing studies on adaptation across divergent light regimes have largely focused on the peripheral sensory systems, often in the context of divergent mate preference [4,5]. However, sensory perception is only the first of many mechanisms within the nervous systems that may experience divergent selection, and mating preferences are only one of many behaviours that may that can be affected by the environment, and consequently contribute to reproductive isolation. Behavioural challenges imposed by novel environments can instead be met by changes in how sensory information is processed, often reflected in differential investment in brain components that refines the sensitivity, acuity to, or integration of, different stimuli.

The intimate relationship between brain structure and ecology is apparent in many comparative studies of neuroanatomy. For example, the expansion of visual pathways in primates [6], cerebellar expansion and refinement of the exterolateral nucleus in electric fish [7–9], the contrasting adaptations of diurnal and nocturnal lifestyles in hawk moths [10], and the independent colonization of cave systems that underlie the radiation in Mexican cavefish [11], all indicate the importance of neuroanatomical adaptations to contrasting ecological needs. However, these comparative studies generally focus on phylogenetically distinct comparisons across relatively distantly related species. At the other extreme, several studies considering inter-specific variation across populations, or between eco-morphs, instead highlight the potential for *plasticity* in brain development to optimize brain structure and function to local conditions [12–14].

Between these population and phylogenetic levels there is a scarcity of information about the role brains play in facilitating speciation across environmental gradients, either through developmental plasticity or the accumulation of heritable changes during ecological divergence. Hence, whether evolutionary changes in neural systems play a causative role in ecological divergence [15], or accumulate later in this process, is unknown. A handful of insect studies have linked specific changes in neural processing to the evolution of reproductive isolation among close relatives, however these specifically focus on divergent host preferences and the detection of host cues [16–22]. Whether brains respond to changes in broader features of the environment, such as luminance or habitat structure, at a similar time scale is yet to be established. Recently, studies of closely related populations on the path to speciation have begun to address this question [11,13,23,24]. Importantly, however, these studies are often unable to disentangle the effects of drift and selection, and have not determined whether hybrids between ecologically distinct populations show disrupted or intermediate brain morphologies that may betray major fitness deficits, and therefore support a more causative role for divergence in neural systems during the incipient stages of speciation.

Here, we investigate the role of heritable divergence in neuroanatomy and gene expression in a clade of closely related *Heliconius* butterflies. *Heliconius* are well known for their bright warning patterns and Müllerian mimicry [25,26]. Speciation events within the *melpomene-cydno* complex are also often associated with ecological transitions [27–29], and habitat partitioning among sister taxa is generally required for complete speciation [30,31]. In particular, within the *melpomene-cydno* clades, coexisting species are often found in “mosaic sympatry”, with sister taxa inhabiting relatively open forest-edge, or closed canopy forest, respectively [30,32,33]. These environmental differences are associated with changes in light environment, and *melpomene/cydno* show evidence of divergence in peripheral eye structure and light sensitivity [34,35]. We hypothesised these differences in habitat-use therefore impose different sensory challenges, leading to consistent, divergent changes in brain structure and function.

## Results and discussion

### Divergence in neuroanatomy in the *Heliconius melpomene-cydno* complex

To investigate the effects of ecological divergence on brain morphology within the *melpomene-cydno* complex, we sampled butterflies from Costa Rica, Panama, Peru, and French Guiana (Figure 1). Where members of the *melpomene* and *cydno* clades are sympatric, the species boundary is maintained by ecological divergence and disruptive selection against hybrids, which now occur at low frequencies [36].

**Figure 1.**
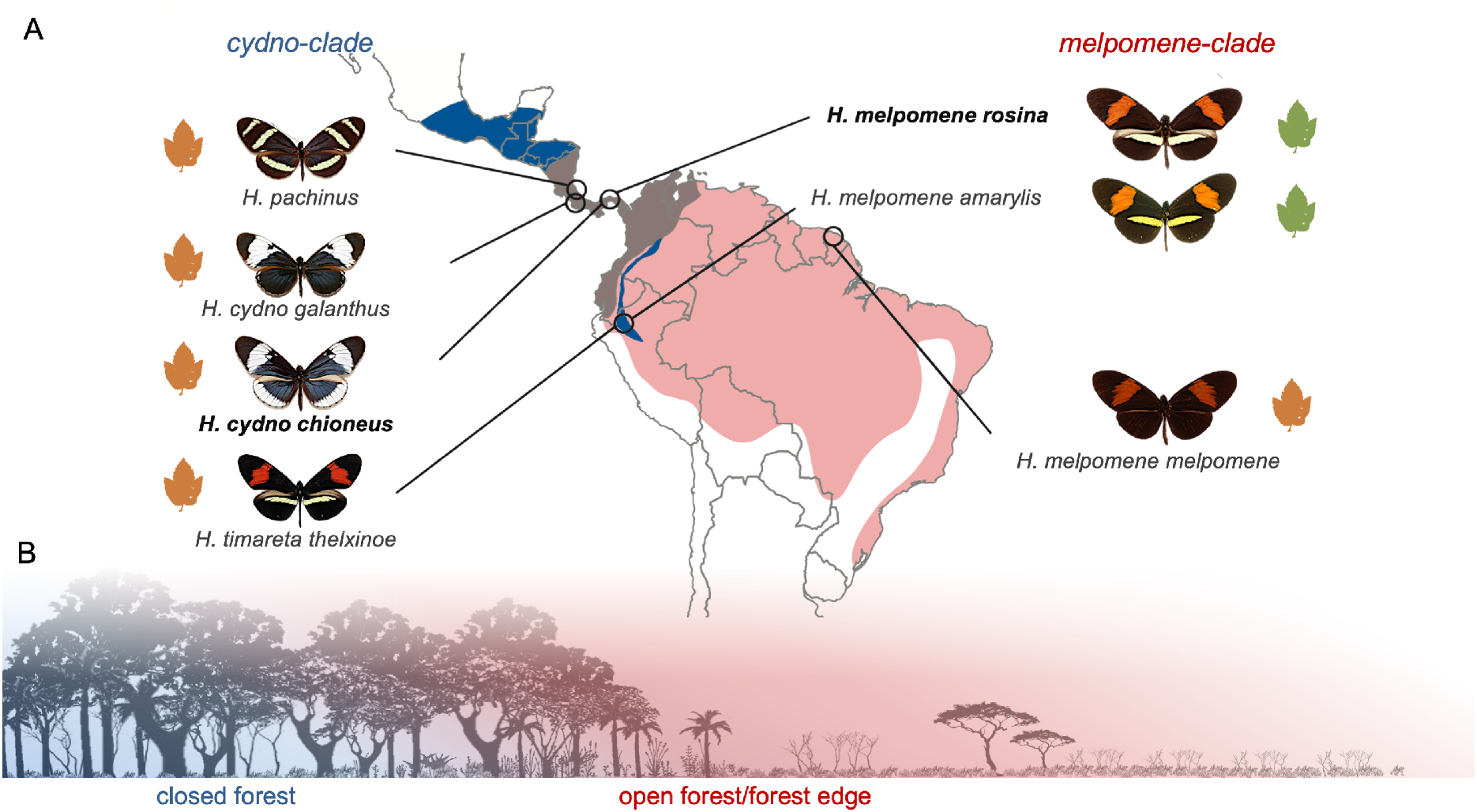
Population sampling and ecological divergence. **(A)** Outline map of Central and South America showing the range of the *cydno* clade (blue), the *melpomene* clade (red), and their overlap (brown). Circles indicate sampled populations in Costa Rica, Panama, Peru and French Guiana with relevant races shown. In the Andes, the *cydno* clade species *H. timareta* is restricted to high elevations, but overlaps with *H. melpomene* at its lower margins. Green *Passiflora sp.* leaves indicate oligophagous races that are host-plant specialists, orange leaves indicate polyphagous host-plant generalists that lay on multiple *Passiflora* species. Races included in the common garden experiments are shown in bold. **(B)** Illustration of niche partitioning between *melpomene* (red; open forest, forest edge) and *cydno* (blue; closed forest).

Across all populations, the average volume of the combined optic lobes neuropils (OL) is significantly larger in *cydno* clade species (including *H. cydno*, *H. pachinus* and *H. timareta)* than *H. melpomene* (n=77, X^2^=17.354, p<0.001), and is not explained by allometric scaling (y-axis shit in OL~rCBR: X^2^=12.260, p<0.001; Figure 2C). Five of the six optic lobe neuropils are significantly larger in the *cydno* clade (Figure 2C,D-I; Table S3), with the sole exception being the lobula. For a given brain size, these neuropils are between 13-27% larger in *cydno*, suggesting altered patterns of investment are unequal across structures. However, in each case the increase is associated with grade-shifts in allometric scaling (Table S4). These structures are vital for summation and parallelization of photoreceptor signals [37–39], and a diverse range of visual processes including colour vision [40–42], shape and motion detection, maneuverability in flight [43,44], and circadian rhythms [45]. The ventral lobula (vLOB), which is only present in some butterflies [46–49], also acts as a relay centre sending visual information to the mushroom body [49], the major site of insect learning and memory.

**Figure 2.**
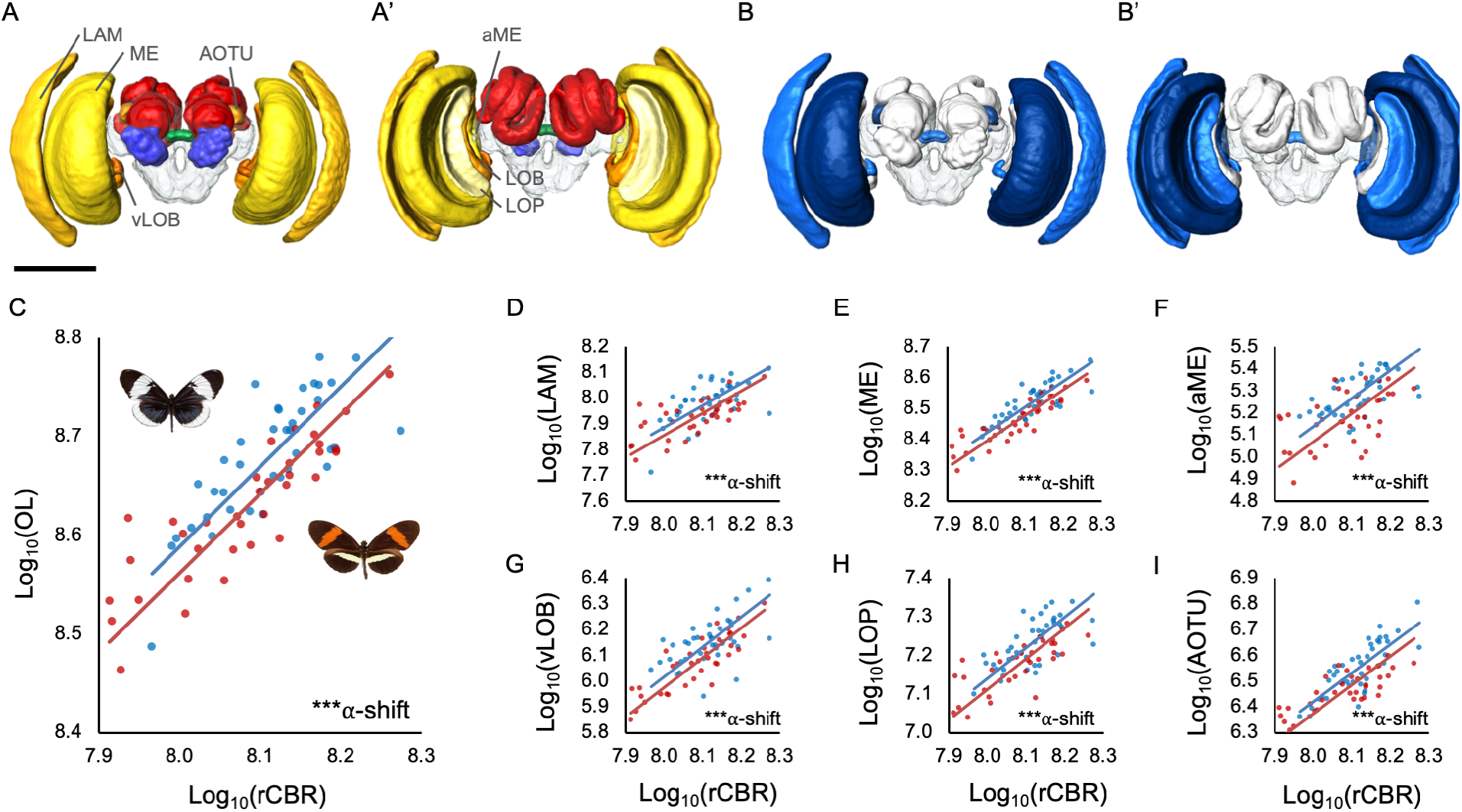
Divergence in brain morphology between *H. melpomene* and *H. cydno*. **(A)** 3D volumetric models of a *Heliconius* brain showing segmented neuropils from anterior (A) and posterior (A’) views; visual neuropils in yellows-oranges, antennal lobe in blue, the central complex in green, the mushroom bodies in red, and the unsegmented rCBR clear. Visual neuropils discussed in the main text are labeled as: LAM is lamina; ME is medulla; aME is accessory medulla; LOB is lobula; LOP is lobula plate; vLOB is ventral lobula; AOTU is anterior optic tubercle. **(B)** 3D volumetric models of a *Heliconius* brain showing segmented neuropils from anterior (A) and posterior (A’) views where blue neuropils are significantly different in size between *H. melpomene* and *H. cydno*, with darker neuropils indicating higher significance. Scale in A/B is 500 μm. **(C)** Grade-shift in the scaling relationship between optic lobe (OL) and central brain (rCBR) volume between *H. melpomene* and *H. cydno*. **(D-I)** Non-allometric shifts in the size of individual visual neuropils between *H. melpomene* and *H. cydno*; lamina (LAM), medulla (ME), accessory medulla (aME), ventral lobula (vLOB), lobula plate (LOP) and anterior optic tubercule (AOTu).

The anterior optic tubercle (AOTU) is also 23% larger in the *cydno*-clade populations (X^2^=10.050, p<0.001). The AOTU is the most prominent optic glomerulus in the central brain, and is involved in processing sky-light and spectral cues, as well as polarised light [50–52]. Contrary to claims that there is a trade-off between investment in major insect visual and olfactory neuropils [53], we find no evidence of volumetric shifts in the antennal lobe (X^2^=0.615, p=0.615). Excluding the AOTU, no other central brain neuropil shows robust evidence for non-allometric expansion (Table S3, S4). Divergence in brain structure is therefore restricted to neuropils associated with visual processing. *H. melpomene* and *H. cydno* occupy forest of different light intensities and physical structure [25,30,32,33], differential investment in these neuropils therefore likely reflects contrasting demands on visual processing. Consistent with this interpretation, *H. cydno* responds to lower intensities of light than *H. melpomene* [34].

### Distinct patterns of intra-clade variation reveal a consistent role of ecology in shaping brain morphology

To further understand the origins of differential investment in visual neuropil, we next considered variation within the *H. cydno* and *H. melpomene* clades. Despite evidence of genetic sub-structuring [54], brain morphology was highly consistent across the four *cydno*-clade populations we sampled, with no neuropil showing significant geographic variation (Table S3B). In contrast, we do find evidence of variation across geographic races of *H. melpomene,* both in total optic lobe volume (X^2^=9.917, p=0.007) and for several of the individual visual neuropils that differentiate the *H. cydno* and *H. melpomene* clades (Table S3B, S4A). These include the largest visual neuropil, the medulla (X^2^=11.161, p=0.004), and the AOTU (X^2^=9.647, p=0.008). Post-hoc analysis reveals that these results are not driven solely by a single divergent population (Table S3C), raising the possibility that *H. melpomene* may occupy more visually heterogeneous habitats than *H. cydno,* and may be tracking local sensory conditions.

Despite greater variability within *H. melpomene*, comparisons between sympatric species pairs suggest a consistent pattern of investment between *melpomene* and *cydno* clade populations. In Panama, *H. m. rosina* and *H. c. chioneus*, are differentiated by total optic lobe volume (X^2^=12.708, p<0.001), with 5 of 7 visual neuropils having larger volumes in *H. cydno* (Table S3A). Similarly, in Peru, *H. m. amaryllis* and *H. timareta thelxinoe* vary in total optic lobe volume (X^2^=6.773, p=0.009) and the two largest visual neuropils, the medulla and lamina (Table S3A). Given *H. m. amaryllis* and *H. t. thelxinoe* are co-mimics and do not appear to distinguish conspecifics using visual cues [55], the shift in visual investment is unlikely to be related to mate choice. In contrast, divergence between *H. c. galanthus* and *H. pachinus*, which are ecologically equivalent but geographically isolated across Costa Rica’s central valley [30,56,57], show no evidence of neuroanatomical divergence despite strong visual mate preferences [57], supporting the causative role of divergent ecologies (Table S4C). Comparisons between *H. m. melpomene*, which is allopatric with respect to *H. cydno*, to all *cydno* populations also detects evidence of divergence in OL volume (X^2^=4.974, p=0.026) with levels of phenotypic divergence comparable to other *melpomene* races (Table S4B). This suggests an absence of strong character displacement for this trait.

### Neuroanatomical differences are heritable

We next reared *H. melpomene rosina* and *H. cydno chioneus* under common garden conditions to determine whether the variation we observe is heritable. As in our comparisons between wild-caught individuals, we observed a non-allometric expansion of the optic lobe in insectary reared *H. cydno* (33%; n=20, X^2^=11.363, p=0.001; Table S5). This was driven by volumetric increases ranging from 24-57% across specific visual neuropils in *H. cydno*, including 5 of the 6 structures that differed between wild-caught individuals (Table S5). The most pronounced shifts were found in the lamina (57% larger, X^2^=13.702, p<0.001), vLOB (49%, X^2^= 6.359, p<0.001) and AOTU (40%, X^2^=21.749, p<0.001). We found no evidence that the extent of divergence for any individual neuropil was higher in wild-caught than common-garden individuals (Table S5C). Differences in brain morphology therefore appear to have a substantial heritable component, and are not the product of environmentally-induced plasticity during development.

### Neuroanatomical divergence is likely driven by natural selection

Allometric scaling among traits, where component sizes vary consistently with total size, is evidence for constraint on trait evolution [58–61], including on brain structure [62,63]. This suggests populations evolving under genetic drift should follow conserved allometric scaling relationships, as is typical among recently diverged taxa [60]. In contrast, our observation of non-allometric variation of brain components, among both wild caught and common-garden reared individuals, strongly implicates divergent natural selection.

To further test the role for selection, we calculated Pst for variation in neuropil volumes in Panamanian *H. m. rosina* and *H. c. chioneus* raised under common garden conditions. Pst is a direct phenotypic analogue of Fst and measures population differentiation relative to the total variance across populations [64]. Comparisons between Pst and Fst can therefore be used as a direct test of selection. After accounting for allometric effects, Pst significantly exceeds genome-wide Fst [65] for total optic lobe size (adjusted-p=0.011), lamina (adjusted-p=0.006), medulla (adjusted-p=0.020), lobula (adjusted-p=0.016), vLOB (adjusted-p=0.005), and AOTU (adjusted-p=0.005), consistent with the action of divergent natural selection. Although inferences made from Pst can be vulnerable to underlying assumptions regarding trait heritability [64], our results are robust across a broad range of quantitative genetic scenarios (Table S7; Supplementary Information).

As a further test for selection acting across the *melpomene-cydno* complex, we performed a partial-Mantel test to assess whether pairwise divergence in brain morphology between wild populations is predicted by levels of neutral genetic divergence (Fst). Here, we expect that the presence of divergent selection would erode the relationship between genetic distance and phenotypic divergence [66]. After allometric correction, only two neuropils, the antennal lobe and lobula, show patterns of divergence consistent with neutral expectations (Table S7). The lack of association for any neuropils with divergent volumes between *H. melpomene* and *H. cydno* again implies our results are not explained by drift.

Together, evidence i) of non-allometric divergence in brain structure, ii) between-species variation that significantly exceed neutral predictions under controlled environmental conditions, and iii) a lack of association between phenotypic and genetic divergence across the *melpomene-cydno* complex, strongly implicates natural selection as the driving force behind the observed differences in neuroanatomical structures.

### Neuroanatomical evolution is mirrored by shifts in neural gene expression

Volumetric changes in neuroanatomy likely indicate difference in cell number or size, which may in turn reflect replicated or divergent circuitry. Shifts in neural physiology or activity are also be behaviorally important but will not be captured in morphometric data. These differences, however, can be captured in differential patterns of gene expression between species. We therefore also examined patterns of gene expression between *H. m. rosina* and *H. c. chioneus,* raised in common garden conditions to control for environmental effects. After accounting for the influence of tissue composition [67], we still detect significant levels of interspecific divergence in expression profiles for age and environment-matched individuals (Figure 4A, Figures S1-3). This pattern is consistent across two independent periods of tissue collection. Differentially expressed genes are enriched for molecular functions linked to cytoskeletal and transmembrane channel activities (Table S8), consistent with changes in brain physiology being achieved through alterations of neuronal wiring or activity.

Differential expression between species could be explained by genetic drift, rather than divergent selection. However, estimated Pst exceeds Fst exceeds genome-wide Fst for 18.5% (305/1647) of differentially expressed genes, strongly implicating divergent selection as a driver behind at least some shifts in neural gene expression. Consistent with this hypothesis, *f_d_*, a measure of shared allelic variation that is used to infer barriers to gene flow [68], is negatively correlated with values of Pst for neural gene expression, even after accounting for variation in recombination rate (X^2^ = 179.0, p ≪ 0.001). Previous genome-wide analyses have highlighted a highly heterogeneous pattern of genetic divergence between *H. m. rosina* and *H. c. chioneus*, with selection against gene flow acting across the genome [65,68]. This suggests the species barrier is determined by multiple, polygenic traits. Because *f_d_*, and by extension Pst, is not clustered across the genome [68], our data is consistent with this inference. We therefore suggest that divergence in neural traits is shaping part of the landscape of genetic differentiation between *H. m. rosina* and *H. c. chioneus*.

### Hybrids show evidence of trait disruption

Reproductive isolation can arise due to a mismatch between intermediate hybrid phenotypes and the environment, such that hybrids suffer lower fitness in either parental environment [1,2]. To explore whether divergent brain structures might contribute to the fitness deficit of hybrids, we produced multiple F1 crosses between *H. m. rosina* and *H. c. chioneus*. We focus on F1 individuals, which account for a major proportion of natural *Heliconius* hybrids [36]. Multivariate analysis of the seven visual neuropils reveals that hybrids show intermediate brain morphologies (Figure 3A; Table S6). This intermediate state is the product of variable dominance effects on specific neuropil (Figure 3B-E; Table S6). Four of the seven neuropil are significantly larger in *H. cydno* than F1 hybrids, but are not significantly different between F1s and *H. melpomene* (Table S6B), suggesting that these are largely influenced by loci with *melpomene-*dominant alleles. In contrast, two neuropil, the lamina and vLOB, are significantly different between F1s and both parental species (Figure 3B,C; Table S6B) implying incomplete or mixed dominance across multiple loci. Importantly, this mosaic pattern also leads to disrupted scaling relationships between some visual neuropil, which may affect the flow and integration of visual information in the brain (Table S6C,D; Figure S1; Supplementary Information).

**Figure 3.**
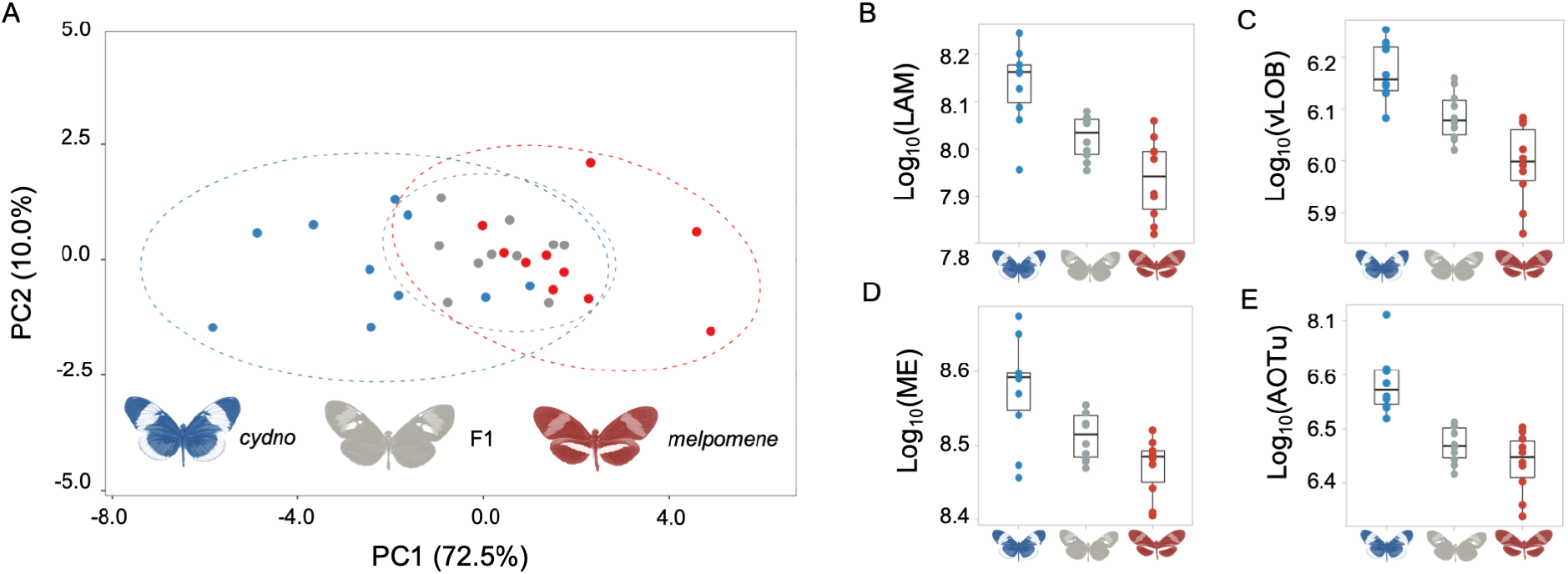
Intermediate brain morphology in *H. m. rosina* x *H. c. chioneus* F1 hybrids. **(A)** Variation in *H. m. rosina* (red), *H. c. chioneus* (blue) and hybrid (grey) brain morphology in a principal component analysis of all segmented neuropils and rCBR. **(B-E)** Examples of neuropils with intermediate volumes in hybrids (B,C), or *melpomene*-like volumes (D, E) in F1 hybrids; lamina (LAM), ventral lobula (vLOB), medulla (ME), and anterior optic tubercule (AOTu).

We observed a similar pattern of hybrid disruption at the molecular level (Figure 4, Figure S3, S4). Focusing on genes that are differentially expressed between *H. cydno* and *H. melpomene*, F1 hybrids cluster outside the range of both parental species (Figure 4A). As was inferred for the visual neuropils, the expression of individual genes show variable patterns of dominance (Figure S5): 36% of differentially expressed genes are ‘*melpomene-*like*’* in F1 hybrids, 21% are ‘*cydno-*like’, and 43% are statistically intermediate. Consistent with divergent selection playing a role in gene expression evolution, genes with intermediate expression in F1 hybrids show increased levels of Pst (X^2^=5825.9, p≪0.001), with a greater proportion (23%) of intermediate genes showing Pst values in excess of genome-wide Fst, compared to genes with *melpomene*-like (9%) or *cydno*-like expression (7%). In contrast, only 0.01% of genes with consistent expression between *H. cydno* and *H. melpomene*, and no genes with transgressive expression in hybrids, show such signatures of selection (Figure 4B). Again, these results are robust across a broad range of quantitative genetic scenarios (Figure S6). In addition, as expected given their enrichment for high Pst values, genes with intermediate hybrid expression are more likely to coincide with regions of reduced gene flow than other differentially expressed genes (X^2^=116.1, p≪0.001; Figure 4C).

**Figure 4.**
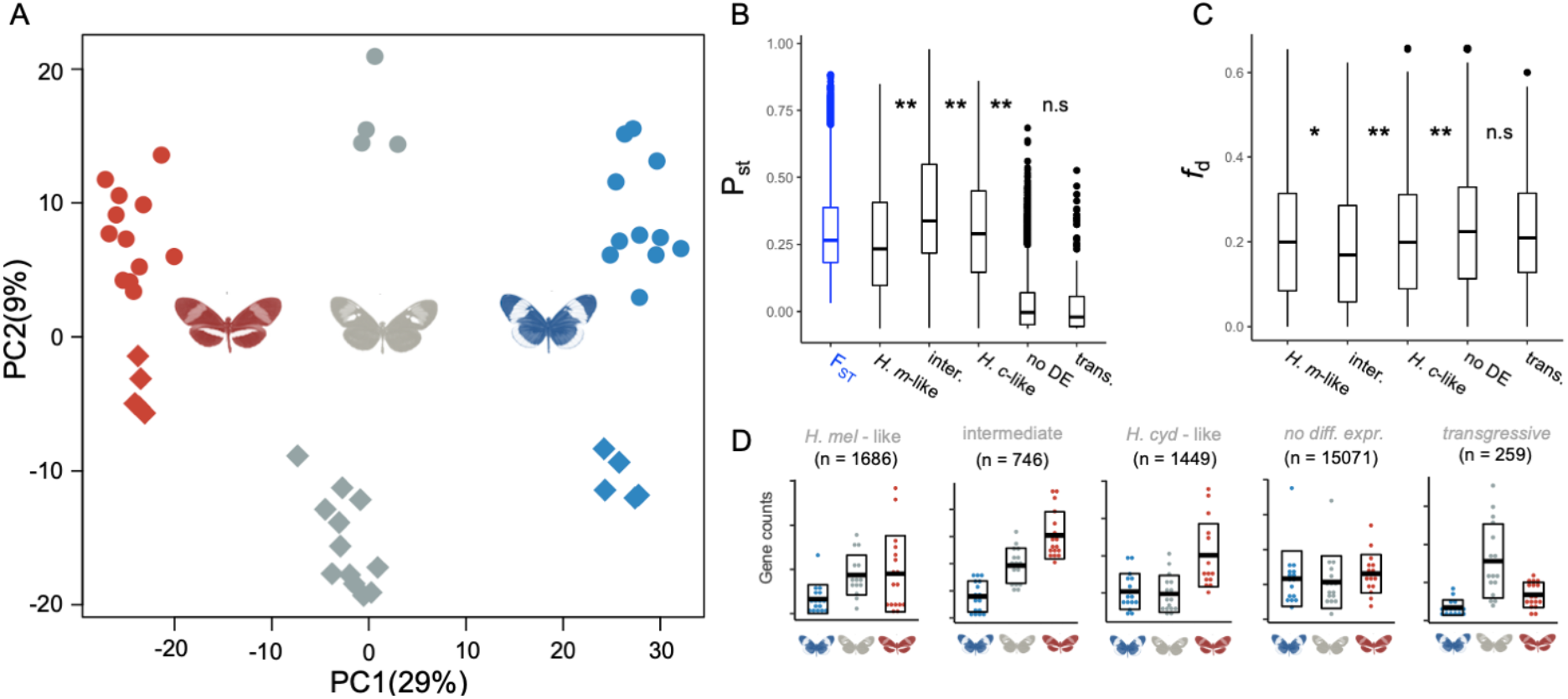
Divergence in gene expression between *H. m. rosina* and *H. c. chioneus*. (A) Principal component analysis of neural gene expression for differentially expressed genes. *H. c. chioneus* samples are colored in blue, F1 hybrids in gray, H. m. rosina in red. Sequencing year is denoted by dot shape: circular (2014), rhomboid (2019). (B) Medians, interquartile ranges and distributions of FST and PST values for genes assigned to different categories based on their expression profiles in F1 hybrids (M-L is *melpomene*-like, I is intermediate, C-L is *cydno*-like, N is no difference, T is transgressive). Below, examples of expression profiles for genes belonging to these different gene categories, with horizontal bars indicating the mean and boxplots delineating +/− sd of the normalized gene counts. n indicates quantity (considering all genes) (C) Median, interquartile range and distributions of admixture proportions (fd), estimated in 100kb windows, between *H. cydno* and *H. melpomene*, for different gene categories. **=p<0.001, *=p<0.01, n.s.=not significant, Kruskal-Wallis test with post-hoc Dunn test, with Bonferroni correction.

Our results therefore reveal both divergence of neural phenotypes between ecologically distinct populations, and disruption of these phenotypes in F1 hybrids. We suggest these intermediate neural phenotypes are likely to act as barriers to gene flow. Divergence in gene expression is a major source of genetic incompatibilities between species [69–71], and cause abnormal development and reduce survival [72]. Hybrid disruption of expression profiles has been reported in diverging species pairs [72–77], however the majority of these studies focus on homogenized whole bodies or gonads. Where organ specific profiles are included, it has been suggested that gonads have an excess of disrupted genes relative to brain tissue [78], and may drive signals from whole-body samples. Nevertheless, some evidence points to the importance of divergence in neural gene during ecological divergence [79,80], and phylogenetic comparisons of neural gene expression in *Heliconius* provide some evidence of selection at deeper time scales [81]. Our data adds clear support for this hypothesis. More broadly, disruption of components of the sensory systems, that co-evolve within species but are under divergent selection between species, likely alters the way in which environmental stimuli are perceived and processed. This occurs at anatomical and molecular levels and may lead to a mismatch between the visual system of hybrids and their sensory environment.

In summary, using a large sample of multiple, geographically disparate populations we have shown that divergent selection during the evolution of micro-habitat partitioning has driven evolution in brain composition and gene expression between *melpomene* and *cydno* clades of *Heliconius* butterflies. These changes are heritable, significantly exceed expected rates of neutral divergence, and result in disrupted traits in F1 hybrids. Neuroanatomical divergence is restricted to the visual neuropils, strongly suggesting that adaptation to contrasting sensory niches contributes to hybrid fitness deficits. This data is consistent with known differences between the two clades in ecology [25,30,32–34,82] and visual sensitivity [34,35]. While disruptive selection on colour pattern has a major role in maintaining reproductive isolation between species [83–85], habitat divergence is thought to be critical to ‘complete’ speciation in *Heliconius* [30,31]. Whether shifts in colour pattern or habitat preference initiate this process is unclear, but given the quality of the aposematic signal is environment and community dependent [86–88], changes in microhabitat preference, and the corresponding neurobiological adaptation to the derived conditions, likely occur at the early stages of divergence. Together, divergent ecological selection on behaviour, and their neural bases, in addition to disruptive selection on mimetic warning patterns would provide strong, coincident barriers to gene flow [89], thereby facilitating speciation.

At a macroevolutionary scale, diverse studies, ranging from recent adaptive radiations in cichlid fish [90] to more ancient diversification of mammals [91], highlight the importance of ecological transitions in driving divergence in sensory regions of the brain. However, whether these changes in brain composition accumulate after ecological transitions, or play a significant role in facilitating them is unclear. Our data provide new evidence that brain evolution has a facultative role in ecological transitions. Our results mirror a previous analysis of divergence in brain morphology between *H. himera* and *H. erato* [23], which are isolated across a steep ecological transition between dense lowland wet forest and more open higher altitude dry forest [28,92]. In this case, heritable shifts in investment are again most notable in sensory neuropils [23]. Similar conclusions can be drawn from the evolution of several fish ecotypes [13,14,93–95], however, here environment-dependent plasticity plays a dominant role in producing population differences [13,14,96]. By demonstrating heritable divergence in brain composition, rates of neural gene expression that exceed neutral expectations, and hybrid disruption at both an anatomical and molecular level, our data provides a robust case for adaptive neural divergence. Given the prevalent role of niche separation and environmental gradients in many adaptive radiations, we suggest that local adaptation in brain and sensory systems may have an underappreciated role during ecological speciation.

## Methods

### Animals

We sampled three pairs of species in Costa Rica, Panama and Peru, and a population of *H. m. melpomene* from French Guiana (Figure 1) with permission from local authorities (Supplementary Information). All wild individuals (n=77) were hand netted and brain tissue was fixed *in situ* in a ZnCl_2_-formalin solution [97] within a few hours of collection. Common garden samples of *H. c. chioneus* and *H. m. rosina* were reared at the Smithsonian Tropical Research Institute’s Gamboa insectaries. Hybrids were produced from multiple *H. c. chioneus* x *H. m. rosina* crosses in 2013 and 2019. Insectary individuals were dissected at 2-3 weeks for neuroanatomical (n=30), and 9-15 days for gene expression samples (n=49) (Table S1,S2).

### Immunohistochemistry and imaging

Brain structure was revealed using immunofluorescence staining against a vesicle-associated protein at presynaptic sites, synapsin (anti-SYNORF1; obtained from the Developmental Studies Hybridoma Bank, University of Iowa, Department of Biological Sciences, Iowa City, IA 52242, USA; RRID: AB_2315424) and Cy2-conjugated affinity-purified polyclonal goat anti-mouse IgG (H+L) antibody (Jackson ImmunoResearch Laboratories, West Grove, PA), obtained from Stratech Scientific Ltd., Newmarket, Suffolk, UK (Jackson ImmunoResearch Cat No. 115-225-146, RRID: AB_2307343). All imaging was performed on a confocal laser-scanning microscope (Leica TCS SP5 or SP8, Leica Microsystem, Mannheim, Germany) using a 10x dry objective with a numerical aperture of 0.4 (Leica Material No. 11506511), a mechanical *z*-step of 2μm and an *x-y* resolution of 512 × 512 pixels. Confocal scans were segmented using Amira 5.5 (Thermo Fisher Scientific) to produce estimates of neuropil volumes.

### RNA extraction and sequencing

Brains were dissected out of the head capsule in cold (4 °C) 0.01M PBS. Total RNA was extracted using TRIzol Reagent (Thermo Fisher, Waltham, MA, USA) and a PureLink RNA Mini Kit, with PureLink DNase digestion on column (Thermo Fisher, Waltham, MA, USA). Illumina 150bp paired-end RNA-seq libraries were prepared and sequenced at Novogene (Hong Kong, China). After trimming adaptor and low-quality bases from raw reads using TrimGalore v.0.4.4 (www.bioinformatics.babraham.ac.uk/projects), Illumina reads were mapped to the *H. melpomene* 2 genome [98]/*H. melpomene* 2.5 annotation [99] using STAR v.2.4.2a in 2-pass mode [100]. We kept only reads that mapped in ‘proper pairs’, using Samtools [101]. The number of reads mapping to each gene was estimated with HTseq v. 0.9.1 (model=union) [102].

### Statistical analyses of neuropil volumes

Non-allometric differences in brain component sizes were estimated using nested linear models in lme4 R [103]. Linear models included each brain component as the dependent variable, the volume of unsegmented central brain neuropil (rCBR), and taxonomic/experimental grouping as independent variables, with sex and country (where relevant) included as random factors. The log-likelihoods of nested models were compared using likelihood ratio tests and a χ^2^ distribution, with sequential Bonferroni correction [104]. For neuropils showing a significant clade/species effect, we subsequently explored the scaling parameters responsible for group differences using SMATR v.3.4-3 [105]. Partial-Mantel tests were performed between pairwise differences in neuropil volumes and Fst [54], controlling for rCBR, using ECODIST [106] with Pearson correlations and 1000 permutations. We calculated Pst using the PSTAT package [107] with a *c*/*h*^*2*^ ratio of 1, and allometric correction with the res() function. The significance of Pst was calculated as the proportion of the Fst distribution [65] that was above each Pst value. Finally, to identify intermediate traits in hybrids we also performed Principal Component Analysis and ANOVAs among parental and hybrid individuals, with post-hoc Tukey-tests to compare group means, using base R packages [108].

### Statistical analyses of gene expression data

Differential gene expression analyses were conducted in DESeq2 [109], including sex and sequencing batch as random factors, with a minimum fold change in expression of 2 to counter effects of tissue composition [67]. We conducted a Principal Component Analysis on rlog-transformed gene count data (as implemented in DESeq2) to inspect clustering of expression profiles. ANOVAs on normalized gene expression counts of species and hybrids, with post-hoc Tukey tests, using base R packages [108]. Pst from normalized gene counts in *H. m. rosina* and *H. c. chioneus* was calculated following Uebbing et al. [110], with *h*^*2*^ set to 0.5 and *c* to 1.0. Estimated admixture proportions (*f*_d_) between *H. m. rosina* and *H. c. chioneus,* and population recombination rates (*rho*) were taken from Martin et al. [68]. To test for an association between low gene flow and high Pst we fitted a linear mixed model: *f*_*d*_ ~ rho + Pst + (*1*|*chromosome*), with a Gaussian distribution, using 100kb non-overlapping windows of *f*_*d*_. GO enrichment tests were performed using InterProScan v.5 [111] to retrieve gene ontology (GO) terms for the Hmel2.5 gene set, and the TopGO package in R [112], using the “elim” algorithm, which corrects for non-independence among GO terms.

Full descriptions of the methodology, and the neuroanatomical dataset are available in the Supplementary Information and will be deposited on DataDryad on manuscript acceptance (DOI pending), along with R code. The raw reads from the genetic dataset will be deposited on The European Nucleotide Archive on manuscript acceptance (accession ID pending).

## Acknowledgements

We are indebted to the environmental agencies in Costa Rica, Panama, Peru, and French Guiana for permissions to carry out this work. We thank Neil Rosser, Ronald Mori Pezo and the Dasmahapatra group for assistance in Peru; the Organization for Tropical Studies at Las Cruces and La Selva, and Le Leona Eco Lodge for assistance in Costa Rica; Adriana Tapia, Moises Abanto, Oscar Paneso, Cruz Batista Saez, Chi-Yun Kuo, Morgan Oberweiser, the McMillan, Jiggins and EBaB labs, and STRI for support at the Gamboa insectaries, Panama. We also thank the University College London Confocal Imaging facility, and Matt Wayland and the Dept. of Zoology Imaging Facility, University of Cambridge, for assistance. This work was funded by a Royal Commission for the Great Exhibition Research Fellowship, a Leverhulme Trust Early Career Fellowship, a short-term STRI Fellowship, British Ecological Society Research Grant (3066) and a NERC IRF (NE/N014936/1) to SHM, and a Deutsche Forschungsgemeischaft Emmy Noether fellowship and research grant (GZ: ME 4845/1-1) to RMM.

## Author contributions

SHM conceived the research with RMM. SHM collected all field and insectary samples for neuroanatomical data, processed and imaged these samples. MR and RMM collected samples for gene expression analyses. SHM and MR analyzed the data. SHM, WOM and RMM secured funding, contributed resources and provided supervision. SHM wrote the manuscript with contributions from all authors.

