## Supplementary Information for "Neural divergence and hybrid disruption between ecologically isolated *Heliconius* butterflies"

### SUPPLEMENTARY INFORMATION, CONTENTS:

1. Materials and methods
   i. Sampling of wild individuals
   ii. Insectary reared individuals
   iii. Neuroanatomy protocols
   iv. Statistical analyses of neuropil volumes
   v. RNA extraction and sequencing
   vi. Statistical analyses of gene expression data

2. Supplementary Results
   i. Tests of neutrality on neuropil volumes
   ii. Hybrid disruption
   iii. Divergence in gene expression
   iv. Classification of gene expression in F1 hybrids
   v. Tests of neutrality on gene expression

3. Supplementary References

4. Supplementary Tables (separate .xls file)
   i. Table S1: Neuroanatomy trait data
   ii. Table S2: Metadata for gene expression samples
   iii. Table S3: Neuroanatomy LMER results for wild caught samples
   iv. Table S4: Neuroanatomy SMATR results for wild caught samples
   v. Table S5: Neuroanatomy LMER and SMATR results for insectary reared samples
   vi. Table S5: Hybrid disruption
   vii. Table S7: Tests of neutrality on neuropil volumes
   viii. Table S8: Differentially expressed genes with categorisation
   ix. Table S9: Gene ontology enrichment

### 1. Materials and methods

#### *1 i. Sampling of wild individuals*

Although they began to diverge ~2million years ago [1], *melpomene* and *cydno* have a long history of persistent gene flow [1,2], and in that sense speciation is considered to be incomplete. Distributed across Central and South America, the species boundary is maintained by ecological divergence and disruptive selection against hybrids [3–6], which now occur at low frequencies [7]. Although deep intra-clade splits are similar in age [1], populations of *melpomene* are currently ascribed to races, while several *cydno* lineages (*timareta, pachinus*) have been promoted to species level [8]. The geographic distribution of the *melpomene* and *cydno* clades allowed us to sample a series of populations, accounting for both geographic divergence on small and continental scales, as well as ecological divergence. All individuals were collected using hand nets and kept alive in glassine envelopes until brain tissue could be fixed within a few hours of collection. Sampling of wild individuals (Table S1) was focused on four countries:

**Panama**

In Panama, *H. c. chioneus* is found in closed forest habitats whereas *H. m. rosina*, occurs in secondary forest [3,9]. We sampled 10 individuals of each species along Pipeline road, Gamboa (elevation 60 m), which transects open to closed forest, and the nearby Soberanía National Park. Samples were collected under permits SEX/A-3-12, SE/A-7-13 and SE/AP-14-18.

**Peru**

*H. timareta* is a member of the *cydno* clade restricted to mid-elevation forest on the eastern Andes. In Peru, *H. t. thelxinoe* is in mosaic sympatry with *H. m. amaryllis*, with which it shares a co-mimetic wing pattern. Like low-elevation *H. cydno*, *H. timareta* is specialised for closed forests [4,10], suggesting micro-habitat partitioning from *H. melpomene* is maintained across the *cydno* clade. Because they are isolated by ecology but not by mimicry ring, this pair provides a ‘control’ for neuroanatomical divergence associated with visual mate cues. 10 individuals of each species were samples in the Escalera region near Tarapoto, Departamento de San Martín (elevation 300-1295 m). Samples were collected under permits 0289-2014-MINAGRI-DGFFS/DGEFFS, 020-014/GRSM/PEHCBM/DMA/ACR-CE, 040–2015/GRSM/PEHCBM/DMA/ACR-CE, granted to Dr Neil Rosser.

**Costa Rica**

*H. c. galanthus* and *H. pachinus* are parapatric species, within the cydno clade, that are restricted to opposite coastal drainages in Costa Rica. In spite of evidence for ongoing gene flow, few hybrids have been collected suggesting that strong reproductive isolation maintains the species barrier [11,12]. Except for differences in colour pattern, there are no known ecological differences between the two taxa suggesting they are ecologically equivalent and the product of allopatric speciation across the Central Valley [11–13]. This provides a 'control' speciation event where we do not expect neuroanatomical divergence between species. 10 *H. c. galanthus* were sampled at La Selva Biological Station (elevation 30-130 m) and Orosí (elevation ~1300 m). 10 *H. pachinus* were sampled at Las Cruces Biological Station (elevation <20 m) and Le Leona eco-lodge on the edge of Corcovado National Park (elevation ~1000 m). A small number of *H. m. rosina* were also collected from these locations. Samples were collected under permit SINAC-SE-GASP-PI-R-2015.

**French Guiana**

At the eastern extreme of its geographic distribution, *H. melpomene* is allopatric with *cydno*. *H. m. melpomene* shares its general ecology with its western relatives, with some exceptions. *H. m. melpomene* is more oligophagus in its larval food plants [14] and there is some suggestion that it uses the forest interior to a greater extent [15], although data supporting this observation is lacking. As we sampled 10 *H. m. melpomene* from forest edge habitats in the Arrondissement of Cayenne (elevation 0-150m), we consider them to have been exposed to similar micro-habitats as *melpomene* in Peru and Panama. We therefore use this population to construct a test of character displacement in brain morphology between sympatric *melpomene/cydno* species. At the time of sampling no permits were required to sample outside National Parks in French Guiana.

***1 ii. Insectary reared animals***

To determine whether any variation we observed was due to environmentally-induced plasticity, we performed common garden experiments focusing on the Panamanian species pair, *H. c. chioneus* and *H. m. rosina*. Insectary-reared individuals were obtained from wild-caught females. Adults were kept under standard conditions in outbred stock cages (c. 1 x 2 x 2 m) of mixed sex and equal densities at the Smithsonian Tropical Research Institute's Gamboa insectaries. These cages are maintained on the edge of the butterfly's native habitat, and light conditions do not substantially deviate from the forest edge environment. Stock cages contained a minimum of 10 females. Because *H. m. rosina* are monophagus, larvae were reared on the species' preferred host plant (*Passiflora menispermifolia* and *P. triloba* respectively). To assess whether hybrid individuals show intermediate or disrupted phenotypes we produced multiple *H. c. chioneus* x *H. m. rosina* crosses in both directions, during two distinct field seasons (2013, 2019). Samples from the 2013 crosses were used for neuroanatomical measurements, while samples from both sets were used to collect gene expression data. We focus on F1 individuals because they are the represent a large portion of hybrids found in natural *Heliconius* hybrids [7] and must survive to produce back crosses with parental species. F1 larvae were reared on *P. triloba.* For both pure species and hybrid crosses, eggs were collected from the host plants on a daily basis over an ~8-week period, and isolated until hatching. Individual larvae were then raised on new growth shoots in outdoor larval cages. After eclosion, adults were aged for 2-3 weeks for the neuroanatomical samples, and 9-15 days for gene expression samples (Table S1, S2). Both sexes are sexually and behaviourally mature at ~8 days [16].

#### *1 iii. Neuroanatomy protocols*

Brains were fixed *in situ* using a $ZnCl_2$-formaldehyde solution, following Ott [17]. Further methodological details and anatomical descriptions of the *Heliconius* brain are available in Montgomery et al. [18]. Briefly, brain structure was revealed using immunofluorescence staining against a vesicle-associated protein at presynaptic sites, synapsin (anti-SYNORF1; obtained from the Developmental Studies Hybridoma Bank, University of Iowa, Department of Biological Sciences, Iowa City, IA 52242, USA; RRID: AB_2315424) and Cy2-conjugated affinity-purified polyclonal goat anti-mouse IgG (H+L) antibody (Jackson ImmunoResearch Laboratories, West Grove, PA), obtained from Stratech Scientific Ltd., Newmarket, Suffolk, UK (Jackson ImmunoResearch Cat No. 115-225-146, RRID: AB_2307343). All imaging was performed on a confocal laser-scanning microscope (Leica TCS SP5 or SP8, Leica Microsystem, Mannheim, Germany) using a 10x dry objective with a numerical aperture of 0.4 (Leica Material No. 11506511), a mechanical *z*-step of 2µm and an *x-y* resolution of 512 x 512 pixels. The *z*-dimension was scaled by 1.52 to correct the artefactual shortening [18]. We assigned image regions to brain components, or neuropils, using the Amira 5.5 (Thermo Fisher Scientific) *labelfield* module and defining outlines based on the brightness of the synapsin immunofluorescence. We reconstructed total central brain volume (CBR), six paired neuropils in the optic lobes (OL), six paired and one unpaired neuropils in the central brain (CBR) in all wild individuals, using the *measure statistics* module to estimate component volumes. In insectary samples the POTu, a small posteriorly located neuropil, was inconsistently stained and was not measured, and in hybrids only neuropils with evidence of divergence between *melpomene* and *cydno* were segmented. The total volume of segmented structures in the CBR was subtracted from total CBR volume to obtain a measure of the remaining, unsegmented CBR (rCBR), which is used as an allometric control throughout. Note that in insectary samples rCBR does not include POTU, and among comparisons including hybrids rCBR is simply CBR minus AOTU volume. Due to the lack of volumetric asymmetry in *Heliconius* neuropils [18] we measured the volume of paired neuropils from one hemisphere, chosen at random unless one hemisphere was damaged, and multiplied the measured volume by two. All volumes were $\log_{10}$-transformed before data analysis.

#### 1 iv. Statistical analyses of neuropil volumes

We identified non-allometric differences between brain component sizes using nested linear models, analysed in the lme4 R package [19]. Linear models included each brain component as the dependent variable, rCBR and taxonomic/experimental grouping as an independent variable, with sex and country (where relevant) included as random factors. The likelihoods of nested models were compared using a $\text{Chi}^2$-test. Correction for multiple testing was performed using a sequential Bonferroni procedure [20]. For neuropils showing a significant clade/species effect, we subsequently explored the scaling parameters responsible for group differences using SMATR v.3.4-3 [21]. Using the standard allometric scaling relationship: $\log y = \beta \log x + \alpha$, where y is the brain component of interest and $x$ is rCBR, we performed tests for significant shifts in the allometric slope ($\beta$) between taxa, followed by two further tests which assume a common slope: 1) for differences in $\alpha$ that suggest discrete 'grade-shifts' in the relationship between two variables, 2) for major axis-shifts along a common slope. Deviation from a shared scaling relationship, by slope or elevation, can indicate an adaptive change in the functional relationship between two brain structures [22].

In addition to our allometrically controlled regressions, we performed two further analyses to explore the role of selection in neuroanatomical divergence. First, using data from wild-caught samples, we performed a Mantel test between pairwise differences in neuropil volumes and two estimates of $F_{ST}$ from Arias et al. [8], based on AFLPs and mtDNA. *H. m. amaryllis* was not included in Arias et al. we therefore use *H. m. malleti* as a surrogate, as these two races are geographically and phylogenetically close [23]. Pairwise differences in neuropil volumes were taken as $\log_{10}(|\text{Population}_A\text{-Population}_B|+1)$. Partial Mantel tests, controlling for pairwise differences in rCBR volumes, were performed using ECODIST [24] with Pearson correlations and 1000 permutations.

Second, with insectary reared samples, we calculated $P_{ST}$ using the PSTAT package [25] initially using a $c/h^2$ ratio of 1, where c is the proportion of the total variance presumed to be due to additive genetic effects across populations, and $h^2$ is the trait heritability. Quantitative genetic parameters for invertebrate neuroanatomy are sorely lacking in the literature, with only a small number of heritability estimates (0.123-0.376) for linear dimensions of *Drosophila* mushroom body size [26]. We therefore also varied the $c/h^2$ ratio assuming $c$ equals 0.25, 0.50, 0.75, 1.00, and $h^2$ equals 0.25, 0.50, 0.75, resulting in ratios of (0.33, 0.67, 1.00, 1.33, 1.50, 2.00, 3.00, 4.00) to test how sensitive the $P_{ST}$ estimates are to these assumptions. $P_{ST}$

calculations were performed on raw, $\log_{10}$-transformed neuropil volumes, and on residual volumes after regressing neuropil volumes against rCBR using the res() function. To test if an individual neuropil's $P_{ST}$ was significantly higher than expected by neutral divergence, we calculated a p-value as the proportion of the $F_{ST}$ distribution [2] (see below) that was above each $P_{ST}$ value, where a $P_{ST}$ value above the 95$^{th}$ percentile of the $F_{ST}$ distribution is taken as evidence of selection.

Finally, we identified intermediate traits in hybrids we also performed Principal Component Analysis and ANOVAs among parental and hybrid individuals, with post-hoc Tukey tests to compare group means, using base R packages [27] (R Core Team, 2013).

#### *1 v. RNA extraction and sequencing*

All samples used for our comparative transcriptomics were reared in common garden conditions (see above). Details of library preparation and sequencing for the 2014 samples are available in Rossi et al. [28]. For all samples, mRNA was extracted from whole brains of age-matched *H. c. chioneus* (n =11), *H. m. rosina* (n = 12), and F1 hybrids in 2014 (n = 4) (see Rossi et al. [28] for further details), and *H. c. chioneus* (n = 5)*, H. m. rosina* (n = 5), and F1 hybrids (n = 12) in 2019. Brains were dissected out of the head capsule in cold (4 ºC) 0.01M PBS solution and include the CBR, OL and ommatidia. For the 2019 samples, total RNA was extracted using aTRIzol Reagent (Thermo Fisher, Waltham, MA, USA), a PureLink RNA Mini Kit, with PureLink DNase digestion on column (Thermo Fisher, Waltham, MA, USA) in 2019. Illumina 150bp paired-end RNA-seq libraries were prepared and sequenced at Novogene (Hong Kong, China). After trimming adaptor and low-quality bases from raw reads using TrimGalore v.0.4.4 (www.bioinformatics.babraham.ac.uk/projects), Illumina reads were mapped to the *H. melpomene* 2 genome [29]/*H. melpomene* 2.5 annotation [30] using STAR v.2.4.2a in 2-pass mode [31]. We kept only reads that mapped in 'proper pairs', using Samtools [32]. The number of reads mapping to each gene was estimated with HTseq v. 0.9.1 (model = union) [33].

#### *1 vi. Statistical analyses of gene expression data*

Differential gene expression analyses between groups were conducted in DESeq2 [34], including sex and sequencing batch as random factors. We considered only those genes showing a 2-fold change in expression level, and at adjusted (false discovery rate 5%) p-values < 0.05, to be differentially expressed due to the potential for tissue composition to drive significant, but low-fold change differences in expression [35].

We conducted a Principal Component Analysis on rlog-transformed gene count data (as implemented in DESeq2) to inspect clustering of expression profiles among groups (species, or hybrids) for all genes, and for differentially expressed genes only.

We performed ANOVAs on normalized gene expression counts of species and hybrids, with post-hoc Tukey tests, and categorized gene expression levels in hybrids as follows:

1. "*melpomene*-like": where F1 vs. *cydno* $p<0.05$ _and_ F1 vs *melpomene* $p>0.05$
2. "*cydno*-like": where F1 vs. *cydno* $p>0.05$ _and_ F1 vs *melpomene* $p<0.05$
3. "intermediate": where F1 vs. *cydno* $p>0.05$ _and_ F1 vs *melpomene* $p>0.05$, **or** *cydno* $p<0.05$ _and_ F1 vs *melpomene* $p<0.05$, with F1s having an intermediate mean expression level between parental species
4. "transgressive": where *cydno* $p<0.05$ _and_ F1 vs *melpomene* $p<0.05$, with F1 having higher or lower mean expression compared to both parental species.

We estimated phenotypic differentiation in gene expression (PST) from normalized gene counts in *H. m. rosina* and *H. c. chioneus*, following Uebbing et al. [36]. For the main analysis we set heritability to be 0.5, but examined the effects of varying heritability in the supplementary results. The distribution of genome-wide genetic differentiation (FST) between *H. m. rosina* and *H. c. chioneus* were retrieved from Martin et al. [2]. To test if an individual gene's PST was significantly higher than expected by neutral divergence, we calculated p-values as the proportion of the FST distribution that was above each PST value. (PST values above the $95^{th}$ percentile of the FST distribution, were considered as showing evidence of selection), We subsequently explored how the frequency of this index of selection varied between gene expression categories (differentially expressed or not, and hybrid categories as defined above) by comparing the proportion of genes showing PST > FST q(95%), in the various gene categories).

To further test whether genes highlighted by these analyses contribute to divergence between *melpomene* and *cydno*, we asked whether genes with PST > FST q(95%), or with intermediate expression in hybrids, were more likely to occur in regions of the genome with low levels of gene flow. We retrieved estimated admixture proportions ($f_d$) between *H. m. rosina* and *H. c. chioneus,* and population recombination rates (*rho*) from Martin et al. [37]. We then investigated the relationship between $f_d$ (estimated in 100kb non-overlapping windows) and PST, accounting for variation in recombination rate, as a way to study whether selection acts against introgression of foreign alleles. In this analysis, we fitted the following generalized linear mixed models (glmm): $f_d$ ~ *rho* + PST + *(1|chromosome),* assuming a Gaussian distribution. We also explored whether genes with intermediate expression in F1s showed higher levels of PST, and lower levels of $f_d$, compared to other genes, with glmm models: *intermediate_Y_or_N ~ X + (1|chromosome), where X =* $f_d$ or PST, assuming a binomial distribution. Inclusion of chromosome as a random factor provides partial correction for the effects of physical linkage between sites, however, we acknowledge this analysis may still be prone to inflated effect sizes due to non-independence of genomic regions. To test for differences in levels of $P_{ST}$ and $f_d$ values among all gene categories we conducted a Kruskal-Wallis test with post-hoc Dunn test (with Bonferroni correction).

Finally, to infer possible overrepresentation of specific molecular functions among gene categories, we first used InterProScan v.5 [38] to retrieve gene ontology (GO) terms associated with every gene annotated in the Hmel2.5 genome. We then conducted a gene set enrichment analysis (Fisher's exact test, $p < 0.01$) with the TopGO package in R [39], using the "elim" algorithm, which corrects for non-independence among GO terms. We identified GO term enrichment tests for differentially expressed genes, genes showing intermediate expression in F1 hybrids, and genes with $P_{ST} > F_{ST}$ q(95%), relative to all other genes, using Fisher's exact tests at $\alpha = 0.05$.

### 2. Supplementary results

#### *2 i. Tests of neutrality on neuropil volumes*

Pairwise volumetric differences in the visual neuropils, or total optic lobe volume, are not generally associated with $F_{ST}$ across populations (Table S7A). Although power is likely limited by the number of populations, the only neuropils that suggest a relationship between trait divergence and $F_{ST}$ are the lobula (LOB), antennal lobe (AL) and both mushroom body components (MBCA, MBLOPE), with only LOB showing an association at $p<0.05$ for $F_{ST}$ estimates from both mtDNA and AFLP data. None of these four neuropils show evidence of non-allometric shifts in relative size between the *melpomene* and *cydno* clades. In contrast, all neuropils that show this pattern of divergence lack associations with $F_{ST}$. Under the association phenotypic drift is linearly associated with neutral genetic divergence this provides evidence for a role of selection in driving divergence in brain composition across the *cydno-melpomene* clade.

$P_{ST}$ estimates based on comparisons between *H. m. rosina* and *H. c. chioneus* reared in common-garden conditions are consistent with this interpretation (Table S7B). With the exception of components of the central complex (PB and CB) all $P_{ST}$ estimates for raw volumes are above the 95$^{th}$ percentile of the $F_{ST}$ distribution, most likely reflecting divergence in total brain size. However, after accounting for allometric variation through regressions against central brain volume (rCBR), significant $P_{ST}>F_{ST}$ effects are only detected for total optic lobe size (OL), lamina (LAM), medulla (ME), lobula (LOB), ventral lobe of the lobula (vLOB), and the anterior optic tubercule (AOTU). All are robust to correcting for multiple tests. Varying the $c/h^2$ ratio suggests these results are widely robust to assumptions about the proportion of genetic variance and heritability (Table S7C). All significant $P_{ST}>F_{ST}$ results are recovered except under low a $c/h^2$ ratio, where the proportion of total variance accounted for by additive genetic variance is low, and heritability is high. A scenario we suspect is unlikely. Even under this scenario, LAM, vLOB and AOTU show significant $P_{ST}>F_{ST}$ results before correcting for multiple tests. Under high $c/h^2$ ratios above 1, the lobula plate (LOP) also has significant $P_{ST}>F_{ST}$.

Taken together at least LAM, ME, vLOB, aME and AOTU show greater degrees of phenotypic divergence between Panamanian *H. m. rosina* and *H. c. chioneus*, <u>and</u> an absence of an association with neutral divergence across the *cydno-melpomene* clade, which is highly suggestive of adaptive evolution, which ultimately affects overall OL size. Our data also suggests that the LOP and LOB have been under divergent selection between *melpomene* and *cydno*, but potentially with less consistency across the clade.

#### *2 ii. Hybrid disruption*

In addition to having intermediate volumes relative to rCBR volume, hybrids also show some evidence of intermediate scaling between pairs of neuropils. Scaling analyses between pairs of visual neuropils in SMATR identify several comparisons with significant deviation in scaling across parental species and F1 hybrids, affecting either the slope or elevation of the scaling relationship (Table S6Di, Dii). Many of these cases reflect pairs of neuropils with direct connections in other insects, including the LOB and vLOB [40], ME and AOTU [41], LOB and AOTU [41], LAM and aME [42], or where there are likely indirect functional connections, e.g. LAM and LOP, which are connected via projections to the MED [43], or the aME and AOTU which both process polarised light [42,44,45].

Post-hoc analyses of tests with p<0.10 suggest that F1 hybrids show potentially intermediate scaling relative to *cydno* and *melpomene*. For example, the elevation constant for scaling between the LAM and aME in F1 hybrids is intermediate between *H. cydno* and *H. melpomene*, although in both cases it is marginally non-significant (*H. cydno* wald = 3.665, p = 0.056; *H. melpomene* wald = 3.758, p = 0.053; Figure S1A). In other cases, scaling in F1 hybrids is significantly different from one parental species, but not both (Table S6D iii; Figure S1B,C). However, different pairs of neuropils show different parental similarities, with some scaling like *H. melpomene* (e.g. aME~AOTU), while other scale like *H. cydno* (e.g. aME~VLOB, LAM~LOB). We suggest that this provides a second potential avenue for hybrid disruption if information flow between multiple neuropils are unbalanced.

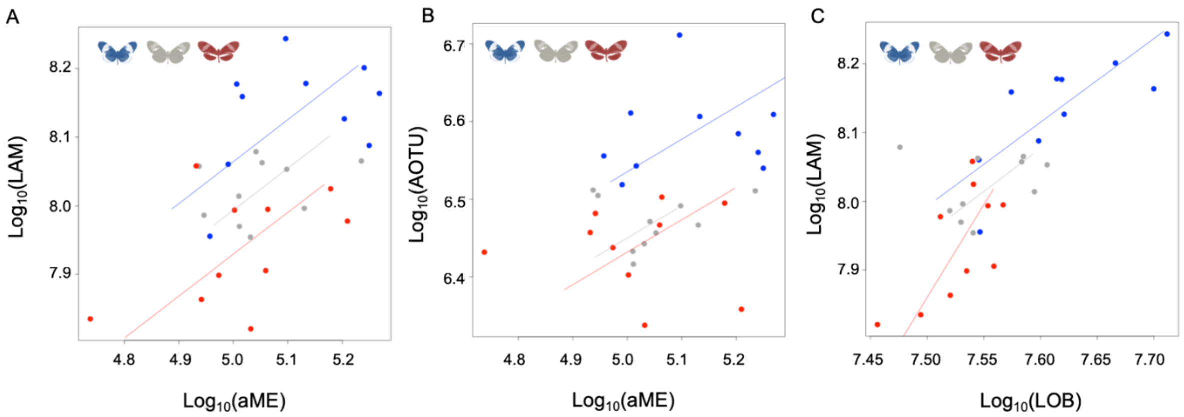


**Figure S1.** Examples of scaling between pairs of neuropils: A) LAM~aME (intermediate), B) AOTU~aME (*melpomene*-like); C) LAM~LOB (*cydno*-like).

### *2 iii. Divergence in gene expression*

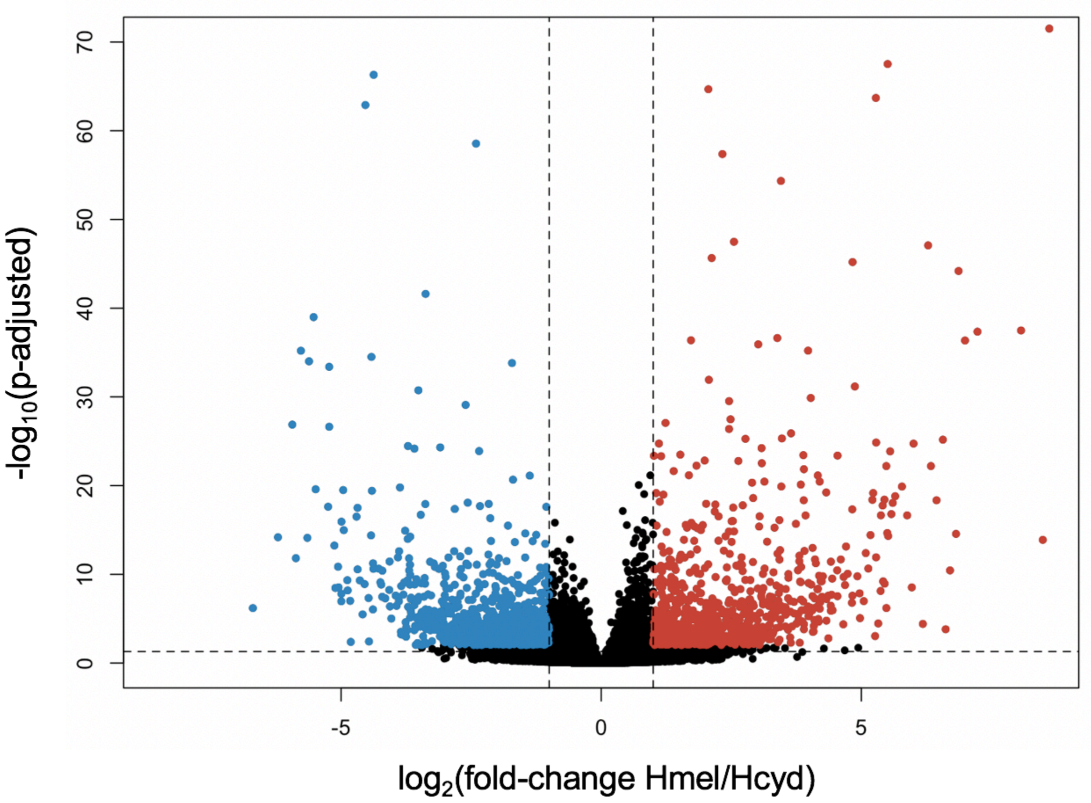


**Figure S2.** Volcano plot for neural gene expression comparisons between *Heliconius melpomene* and *H. cydno*. Vertical dotted lines indicate the thresholds of a 2-fold change in expression (at x values of -1 and 1), the horizontal dotted line indicates significance (p-adjusted<0.05) in the test for differential expression (as conducted in DESeq2). Differentially expressed genes are colored in blue if up-regulated in *cydno*, and red if up-regulated in *melpomene*. Note that 3 outliers (with very low associated p-values) were removed for clarity.

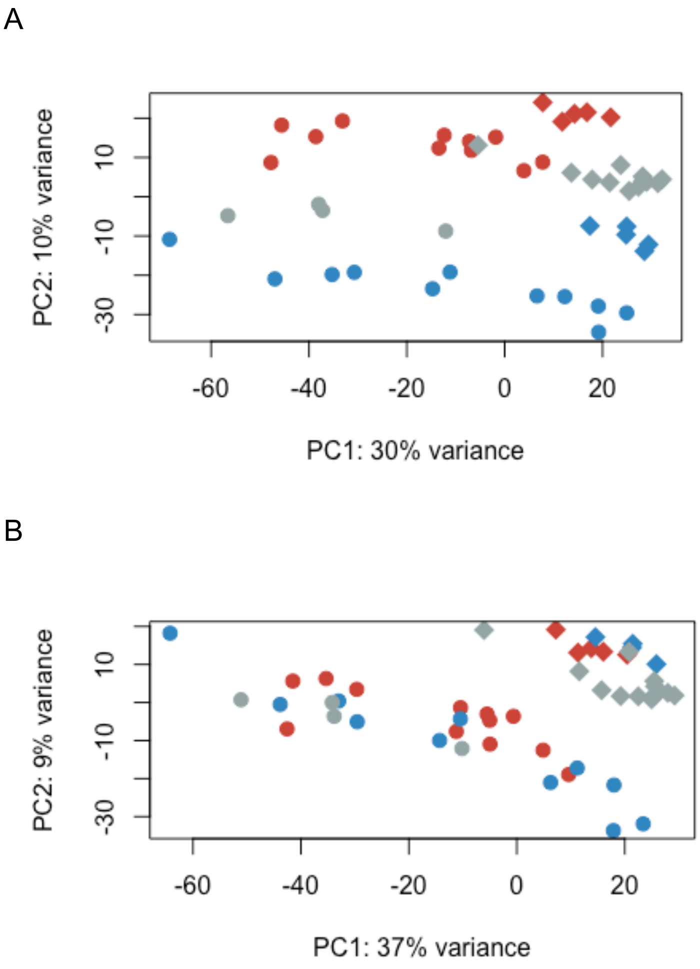


**Figure S3.** Principal component analyses of expression level profiles of (**A**) all genes and of (**B**) genes that were not detected as differentially expressed (see also, Figure 4). *H. cydno* samples are colored in blue, F1 hybrids in gray, *H. melpomene* in red. Hybrids show reduced intermediary in gene expression level when taking all genes into account, and are not intermediate when non-differentially expressed genes are analysed. However, a trend for dominance of the *melpomene* alleles is also evident across all genes (of all genes: 8.7% are *melpomene-like*, 7.5% *cydno*-like, 3.9% statistically intermediate, 1.3% transgressive, 78.3% show no difference between species), Sequencing batch is denoted by the dot shape: circular (batch 2014) and rhomboid (batch 2019).

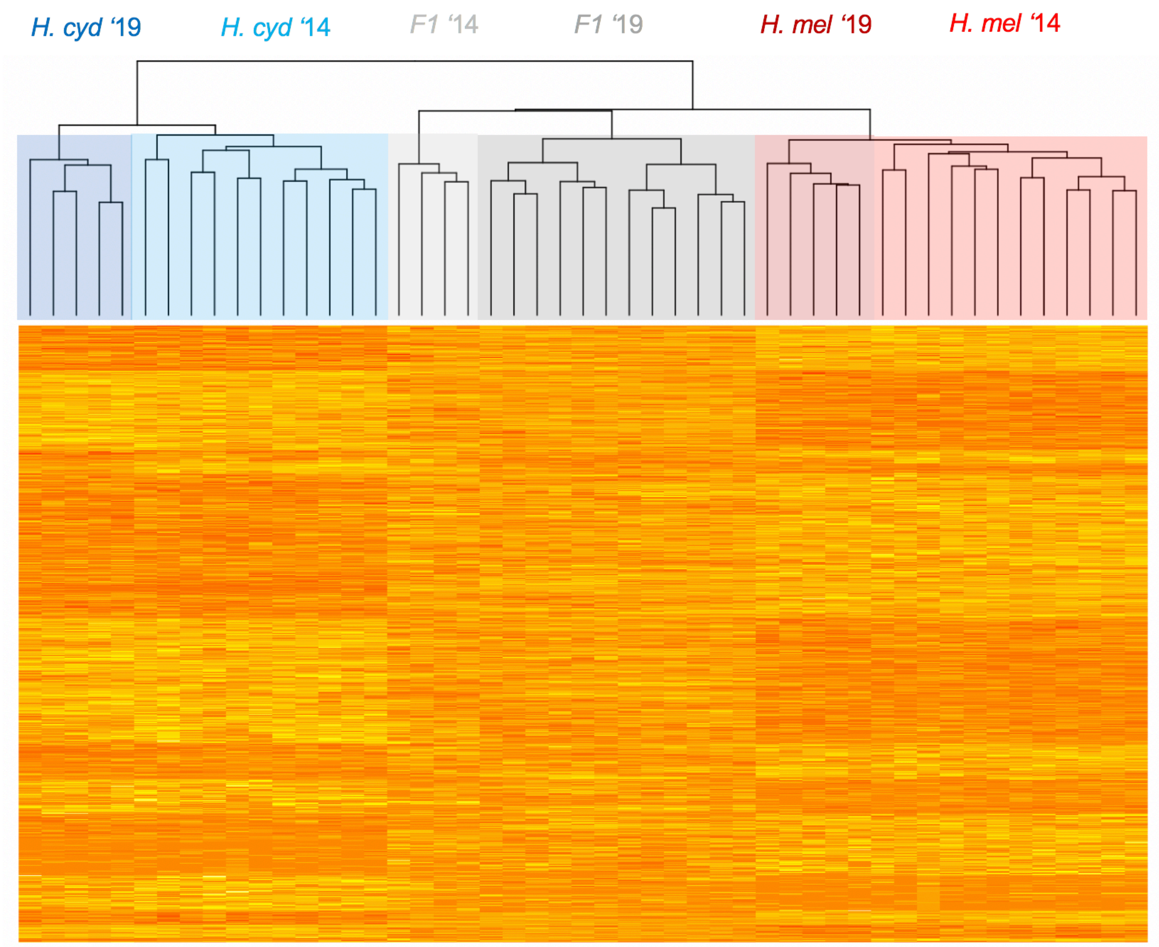


**Figure S4.** Dendogram of neural expression profiles for *H. cydno*, F1 hybrids and *H. melpomene,* for genes detected to be differentially expressed between *cydno* and *melpomene* (different sequencing batches, 2014 and 2019, are highlighted with different shades of blue *for H. cydno,* gray for F1s, and red for *H. melpomene*).

#### *2 iv. Classification of gene expression in F1 hybrids*

Our classification of gene expression patterns in hybrids relative to both parental species provides insights into the disruptive nature of hybridisation on expression profiles. Figure S5 provides illustrative examples of the pattern of variance characteristic of each gene category. Among differentially expressed genes, 589 were *melpomene*-like, 344 were *cydno*-like, 12 were ‘transgressive’, and 701 were intermediate between parental distributions. Considering all genes regardless of their differential expression between species, 1686 were classed as *melpomene*-like, 1449 were *cydno*-like, 259 were ‘transgressive’, and 746 were intermediate

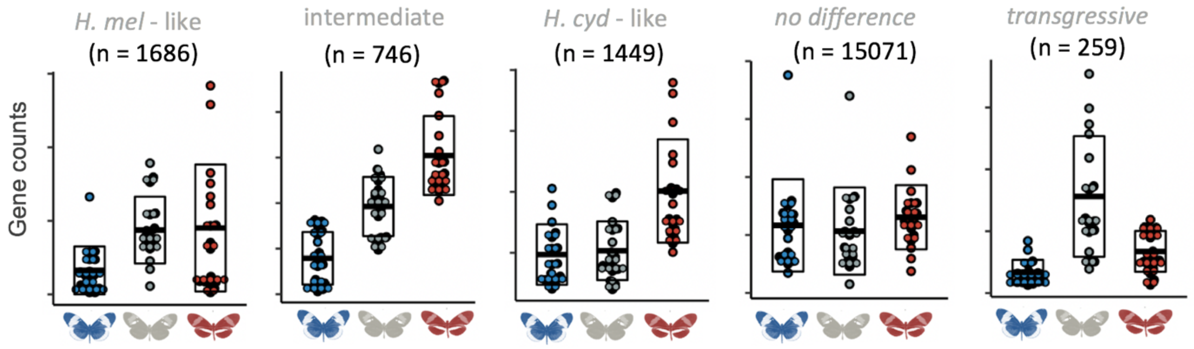


**Figure S5.** Example of expression profiles for genes assigned to the different categories mentioned (e.g. intermediate in F1s). “n =” denotes the number of genes classified in each category. y-axis indicates the (rlog) normalized gene count. Dots correspond to individual samples, and are colored in blue for *H. cydno* samples, in gray for F1 hybrids, and in red for *H. melpomene.* Horizontal black bars indicate mean, with boxplots delineating + and - one standard deviation, of normalized gene counts.

### *2 v. Tests of neutrality on gene expression*

PST estimates vary significantly between gene categories (Kruskal-Wallis test with post-hoc Dunn test with Bonferroni correction, p<0.001 in pairwise comparisons except between transgressive and “no differentially expressed” genes), with 11% (n=432/3881) of genes showing intermediate or species-like expression in hybrids having PST estimates within the 95$^{th}$ percentile of the FST distribution, compared to 0.02% (n=3/15330) for non-differentially expressed genes. The proportion of genes with significant PST estimates is highest within genes with intermediate expression in hybrids 23% (n=169/746), followed by *melpomene* and *cydno*-like genes (9% (n=159/1686) and 7% (n=104/1449) respectively), and lowest within transgressive genes (0% (n=0/259)). To test how these results varied under different assumed heritabilities, we recalculated PST under five $h^2$ values (Figure S5). As expected, the results are consistent across this range, with increased support for selection at lower $h^2$ values.

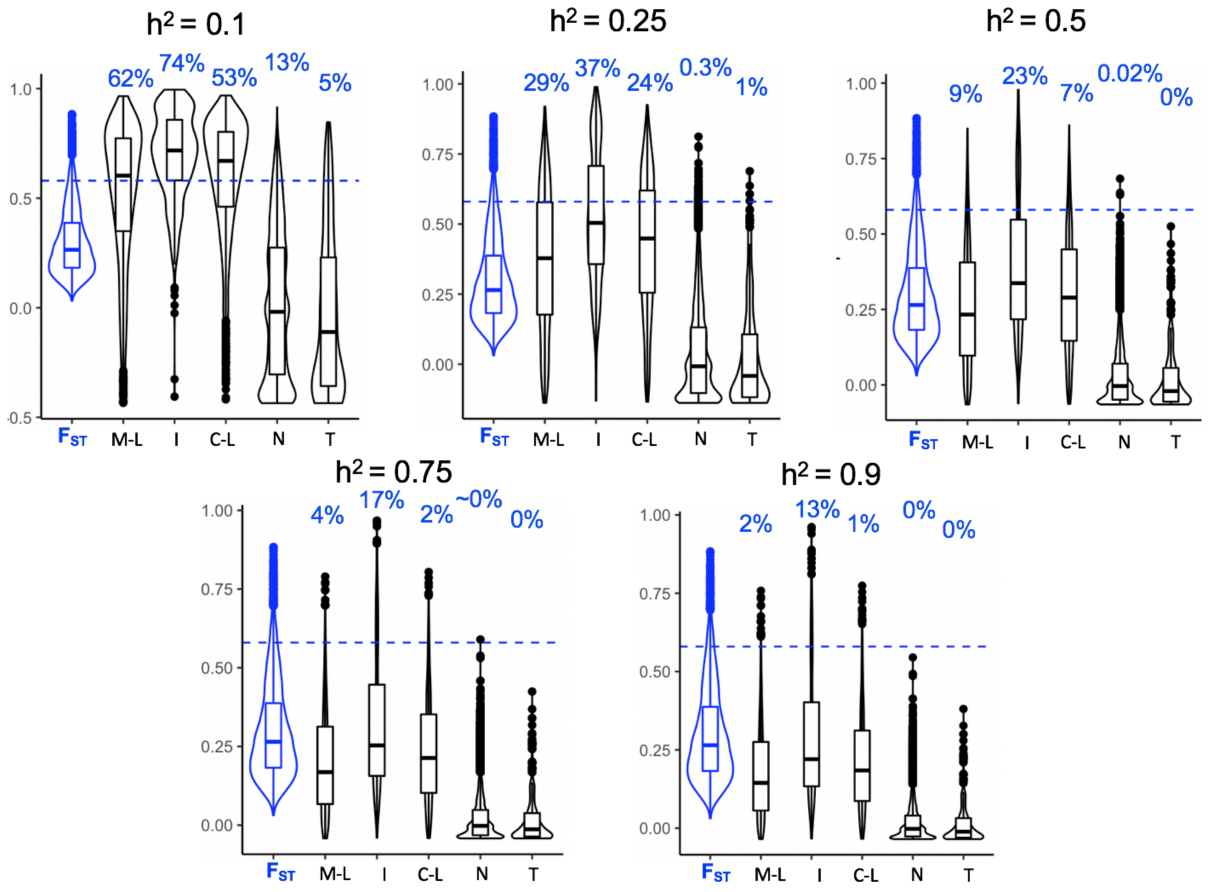


**Figure S6.** Median, interquartile range and distributions of FST and PST values (for different gene categories), where PST was estimated with varying levels of heritability ($h^2$), indicated on top of each panel. Percentages (in blue) indicate the percentages of genes with PST value higher than the 95% quantile of FST (indicated by a horizontal dotted blue line), in each category.
